# SpoIVA-SipL complex formation is essential for *Clostridioides difficile* spore assembly

**DOI:** 10.1101/522235

**Authors:** Megan H. Touchette, Hector Benito de la Puebla, Priyanka Ravichandran, Aimee Shen

**Author notes:** Current address: Nova Biomedical, Waltham, MA, USA. Address correspondence to, Aimee Shen.

## Abstract

Spores are the major infectious particle of the Gram-positive nosocomial pathogen, *Clostridioides* (formerly *Clostridium*) *difficile*, but the molecular details of how this organism forms these metabolically dormant cells remain poorly characterized. The composition of the spore coat in *C. difficile* differs markedly from that defined in the well-studied organism, *Bacillus subtilis*, with only 25% of the ~70 spore coat proteins being conserved between the two organisms, and only 2 of 9 coat assembly (morphogenetic) proteins defined in *B. subtilis* having homologs in *C. difficile.* We previously identified SipL as a clostridia-specific coat protein essential for functional spore formation. Heterologous expression analyses in *E. coli* revealed that SipL directly interacts with *C. difficile* SpoIVA, a coat morphogenetic protein conserved in all spore-forming organisms, through SipL’s C-terminal LysM domain. In this study, we show that SpoIVA-SipL binding is essential for *C. difficile* spore formation and identify specific residues within the LysM domain that stabilize this interaction. Fluorescence microscopy analyses indicate that binding of SipL’s LysM domain to SpoIVA is required for SipL to localize to the forespore, while SpoIVA requires SipL to promote encasement of SpoIVA around the forespore. Since we also show that clostridial LysM domains are functionally interchangeable at least in *C. difficile*, the basic mechanism for SipL-dependent assembly of clostridial spore coats may be conserved.

**Importance:** The metabolically dormant spore-form of the major nosocomial pathogen, *Clostridioides difficile*, is its major infectious particle. However, the mechanisms controlling the formation of these resistant cell types are not well understood, particularly with respect to its outermost layer, the spore coat. We previously identified two spore morphogenetic proteins in *C. difficile*: SpoIVA, which is conserved in all spore-forming organisms, and SipL, which is conserved only in the Clostridia. Both SpoIVA and SipL are essential for heat-resistant spore formation and directly interact through SipL’s C-terminal LysM domain. In this study, we demonstrate that the LysM domain is critical for SipL and SpoIVA function, likely by helping recruit SipL to the forespore during spore morphogenesis. We further identified residues within the LysM domain that are important for binding SpoIVA and thus functional spore formation. These findings provide important insight into the molecular mechanisms controlling the assembly of infectious *C. difficile* spores.

## Introduction

The Gram-positive pathogen *Clostridioides* (formerly *Clostridium*) *difficile* is a leading cause of antibiotic-associated diarrhea and gastroenteritis in the developed world (1, 2). Since *C. difficile* is an obligate anaerobe, its major infectious particle is its aerotolerant, metabolically dormant spore form (3, 4). *C. difficile* spores in the environment are ingested by susceptible hosts and transit through the gastrointestinal tract until they sense specific bile salts in the small intestine that trigger spore germination (5). The germinating spores outgrow into vegetative cells in the large intestine, which then produce the glucosylating toxins responsible for disease symptoms (6). A subset of these vegetative cells will initiate the developmental program of sporulation, producing the infectious spores needed for this organism to survive exit from the host (7, 8).

The basic architecture of spores is conserved across endospore-forming bacteria: a central core consisting of partially dehydrated cytosol is surrounded by a protective layer of modified peptidoglycan called the cortex, which is in turn encased by a series of proteinaceous shells known as the coat (9). The cortex is critical for maintaining spore dormancy and conferring resistance to heat and ethanol, while the coat acts as a molecular sieve that protects the spore from enzymatic and oxidative insults (9-11). The cortex is assembled on top of the thin layer of vegetative cell wall that is sandwiched in between two membranes known as the inner forespore and outer forespore membranes. The outer forespore membrane derives from the mother cell and encases the developing forespore during engulfment by the mother cell. During this time, a series of self-polymerizing proteins will assemble on the outer forespore-membrane and eventually form the concentric layers of protein that define the spore coat (10).

The mechanisms controlling coat assembly have been studied for decades in *Bacillus subtilis*, where the key coat morphogenetic proteins that control the recruitment and assembly of the coat layers have been identified (10, 12). The innermost layer, known as the basement layer, is formed through the coordinated actions of SpoVM, SpoIVA, and SpoVID. SpoVM is a small amphipathic helix that embeds itself in the forespore membrane (13) and directly interacts with SpoIVA (14), facilitating SpoIVA’s assembly around the forespore (14). SpoIVA is a self-polymerizing ATPase (15) that binds SpoVM through residues in SpoIVA’s C-terminal region (14). In the absence of SpoVM, SpoIVA forms a single focus on the forespore and fails to encase the forespore (14, 16). SpoIVA also recruits SpoVID to the forespore (17), and their interaction promotes the encasement of both proteins around the forespore (17).

Loss of any one of these coat morphogenetic proteins in *B. subtilis* prevents recruitment of additional coat proteins to the forespore and causes the polymerized coat to mislocalize to the mother cell cytosol, at least in the case of *spoIVA* (18) and *spoVID* (19) mutants. Loss of either SpoIVA or SpoVM prevents cortex assembly and thus heat-resistant spore formation (~10^−8^ defect) due to activation of a quality control pathway conserved in the Bacilli (18, 20, 21). In contrast, loss of SpoVID results in only an ~10-fold defect in heat-resistant spore formation but an ~1000-fold decrease in lysozyme resistance, consistent with the presence of the cortex layer in a *spoVID* mutant (19). It should be noted that a more recent study observed that the lysozyme sensitivity of a clean *spoVID* deletion mutant is less severe than that of a transposon mutant (~10-fold vs. ~1000-fold) (22).

Interestingly, while SpoIVA and SpoVM appear to be conserved in all spore-forming organisms, SpoVID is conserved only in the Bacilli (23). Of the 9 coat morphogenetic proteins that have been defined in *B. subtilis*, only two have homologs in *C. difficile*, namely SpoIVA and SpoVM (24). While we previously showed that SpoIVA is critical for spore formation in *C. difficile* (25), we surprisingly found that SpoVM is largely dispensable for functional *C. difficile* spore formation, despite abnormalities in coat adherence to the forespore being observed (26). Furthermore, unlike *B. subtilis*, both *spoVM* and *spoIVA* mutants in *C. difficile* produce cortex (25, 26), although some abnormalities in cortex thickness are observed (26).

Although SpoVID is not conserved in the Clostridia, we previously showed that *C. difficile* produces a functional homolog to SpoVID called SipL (CD3567). Similar to *B. subtilis* SpoVID (17), *C. difficile* SipL is required for proper localization of the coat around the forespore (25), and SipL directly interacts with *C. difficile* SpoIVA (25), even though SipL and SpoVID exhibit no sequence homology outside of their shared C-terminal LysM domain (25) (**Fig. 1A**). However, unlike *B. subtilis* SpoVID, *C. difficile*’s LysM domain directly binds to SpoIVA (25), whereas SpoVID’s LysM domain is dispensable for SpoVID binding to SpoIVA (17). Furthermore, loss of *C. difficile* SipL causes a severe heat-resistance defect (<10^6^) (25), in contrast with the mild defects observed in *B. subtilis spoVID* mutants (19, 22).

**Figure 1.**
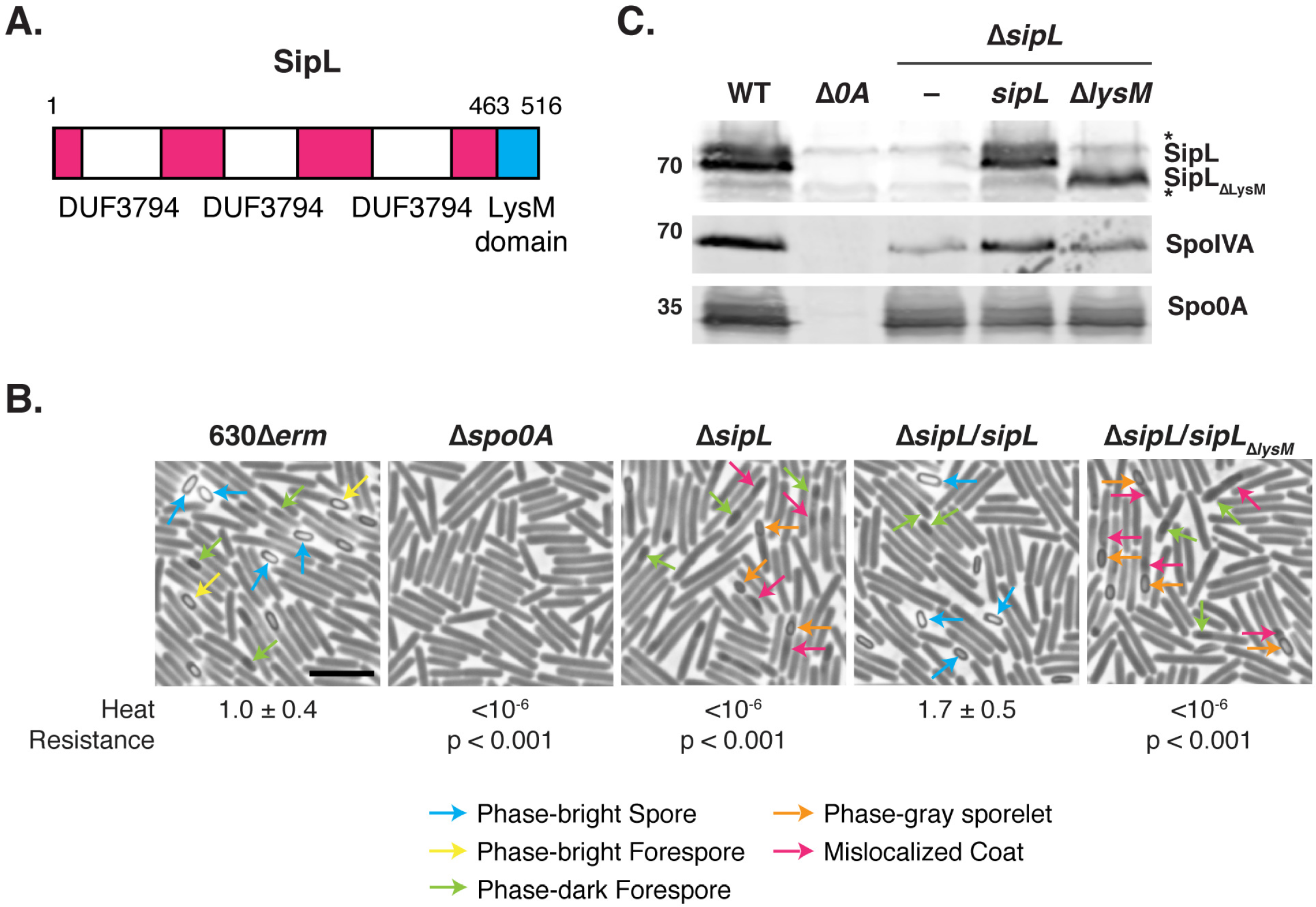
The LysM domain of* C. difficile* is essential for spore formation. (A) Schematic of *C. difficile* SipL domain structure. SipL contains three domains of unknown function (DUF3794) and a C-terminal LysM domain. (B) Phase-contrast microscopy analyses of the indicated *C. difficile* strains ~20 hrs after sporulation induction. Δ*sipL* was complemented with either the wild-type allele or one encoding a deletion of the LysM domain (Δ*sipL*/*sipL_ΔlysM_*). Arrows mark examples of sporulating cells at different stages of maturation: blue arrows highlight phase-bright free spores; yellow arrows mark mature phase-bright forespores, which are formed following cortex formation (22, 24); green arrows highlight immature phase-dark forespores; orange arrows highlight phase-gray sporelets, which produce a phase-dark ring surrounding the forespore but do not become phase-bright; and pink arrows demarcate regions suspected to be mislocalized coat based on previous studies (26, 30). Heat resistance efficiencies were determined from 20-24 hr sporulating cultures and represent the mean and standard deviation for a given strain relative to wild type based on a minimum of three independent biological replicates. Statistical significance relative to wild type was determined using a one-way ANOVA and Tukey’s test. Scale bar represents 5 μm. The limit of detection of the assay is 10^−6^. (C) Western blot analyses of SipL, SpoIVA, and Spo0A. SipL was detected using an antibody raised against SipL lacking its LysM domain (i.e. SipL_ΔLysM_). Asterisks indicate non-specific bands detected by the SipL_ΔLysM_ antibody. SpoIVA levels were analyzed because of the prior finding that SpoIVA levels are reduced in the absence of SipL (25). Modest decreases in SipL levels were observed in the Δ*sipL* and Δ*sipL*/*sipL*_Δ*lysM*_ strains. Spo0A was used as a proxy for measuring sporulation induction (4, 25). The western blots shown are representative of the results of three independent biological replicates.

While we previously showed that SipL binds to SpoIVA through its LysM domain using a heterologous *E. coli* expression system (25), in this study we tested the hypothesis that this interaction is critical for *C. difficile* spore formation using deletion and co-immunoprecipitation analyses in *C. difficile*. We also identified residues in the LysM domain important for both SipL function and binding to SpoIVA. Lastly, we determined the requirement for the LysM domain to localize SipL to the forespore and the localization dependencies of SpoIVA and SipL

## RESULTS

### The LysM domain is required for SipL function

Based on our prior finding that SipL binding to SpoIVA depends on SipL’s LysM domain when heterologously produced in *E. coli* (25), we sought to test whether SipL binding to SpoIVA is critical for SpoIVA and/or SipL function during *C. difficile* sporulation. To this end, we expressed a construct encoding a deletion of the LysM domain in a Δ*sipL* strain we previously constructed (27) to generate strain Δ*sipL/sipL*_Δ*lysM*_. As with all constructs tested in this manuscript, the *sipL*_Δ*lysM*_ construct was expressed from the native *sipL* promoter from the ectopic *pyrE* locus of 630Δ*erm* background using the *pyrE*-based allele coupled exchange system (28). Functional spore formation in the *sipL*_Δ*lysM*_ and wild-type *sipL* complementation strains was then assessed using a heat resistance assay. In this assay, sporulating cultures are heat-treated to kill vegetative cells, while heat-resistant spores in the cultures capable of germinating and forming colonies on plates are enumerated and compared to colony-forming unit (CFU) counts from untreated samples. The LysM domain mutant (Δ*lysM*) was defective in heat-resistant spore formation at levels similar to the parental Δ*sipL* strain (~6-log decrease, **Fig. 1B**). In contrast, the wild-type *sipL* construct expressed from the *pyrE* locus (Δ*sipL*/*sipL*) fully complemented the parental Δ*sipL* strain.

To ensure that the inability of the *sipL*_Δ*lysM*_ construct to complement the Δ*sipL* strain was not due to destabilization of SipL_ΔLysM_, we analyzed SipL levels in the different *sipL* complementation strains using an antibody raised against SipL lacking its LysM domain. These analyses revealed that loss of the LysM domain did not reduce SipL levels in sporulating cells relative to wild-type or the wild-type *sipL* complementation strain (**Fig. 1C**). Thus, the sporulation defect of the Δ*sipL*/*sipL*_Δ*lysM*_ strain is because SipL lacking its LysM domain is non-functional. Consistent with our prior finding that SipL helps stabilize SpoIVA (25), SpoIVA levels were slightly reduced in the Δ*sipL* and Δ*sipL*/*sipL*_Δ*lysM*_ strains. However, the reduction in SpoIVA levels did not appear as large or as consistent in our analyses of the 630Δ*erm* Δ*sipL* mutants relative to the previously characterized JIR8094 *sipL*::*erm* Targetron mutant (25), which may reflect strain-specific differences between JIR8094 and 630Δ*erm* (29).

Phase-contrast microscopy analyses revealed that the Δ*lysM* strain resembled the Δ*sipL* strain in that it failed to produce phase-bright spores (**Fig. 1B**). Instead, these defective strains occasionally produced phase-gray sporelets (orange arrows) (20) that did not achieve the oval shape and phase-bright contrast of wild-type spores (**Fig. 1B**). Furthermore, phase-dark regions (pink arrows) were visible in the mother cell cytosol of Δ*lysM* and Δ*sipL* strains unlike wild type and the wild-type complementation strain. These regions likely correspond to mislocalized spore coat based on prior work (25, 26, 30). To address this possibility, we visualized the spore coat of these strains using transmission electron microscopy (TEM). Δ*sipL/sipL*_Δ*lysM*_ and Δ*sipL* cells failed to localize coat around the forespore in analyses of >50 sporulating cells. Instead, polymerized coat detached from the forespore and mislocalized to the mother cell cytosol in ~40% of these cells (pink arrow, **Fig. 2**), while polymerized coat appeared to slough off the forespore (previously termed “bearding,” (26) yellow arrow, **Fig. 2**) in ~40% of these strains. Neither of these phenotypes was detected in wild type or the wild-type *sipL* complementation strain, where coat was localized around the forespore in ~95% of wild-type and Δ*sipL*/*sipL* cells or otherwise was not yet visible. Taken together, these analyses indicate that loss of SipL’s LysM domain results in coat mislocalization and impairs its adherence to the forespore.

**Figure 2.**
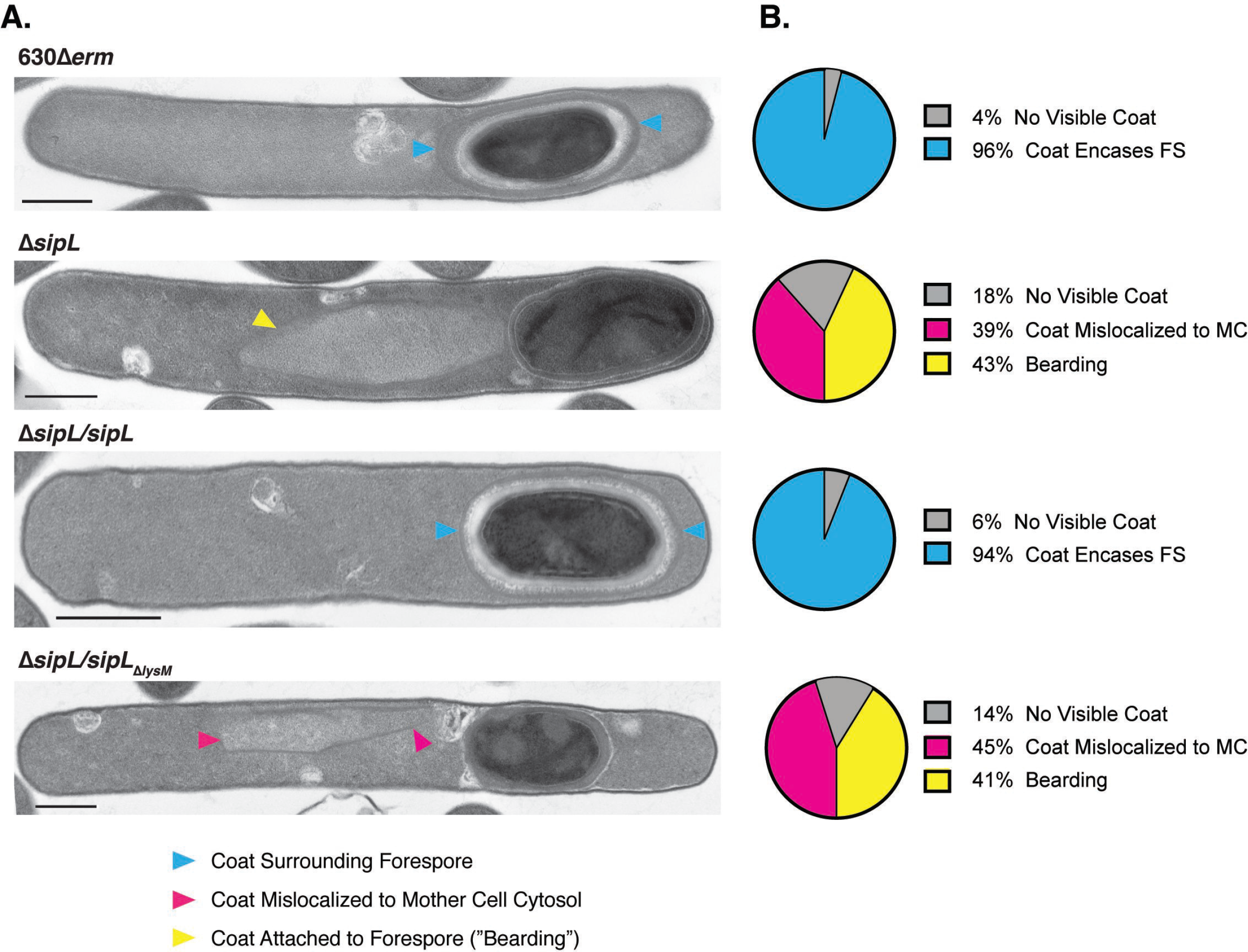
Loss of SipL’s LysM domain results in coat mislocalization defects. (A) Transmission electron microscopy (TEM) analyses of wild-type 630Δ*erm,* Δ*sipL*, and Δ*sipL* complemented with either wild-type *sipL* or *sipL* encoding a LysM deletion (*sipL*_Δ*lysM*_) after 23 hrs of sporulation induction. Scale bars represent 500 nm. Blue arrows mark properly localized coat, i.e. surrounding the entire forespore (FS), whereas pink arrows mark coat that has completely detached from the forespore and is found exclusively in the mother cell (MC) cytosol. Yellow arrows mark cells where coat appears to be detaching from the forespore but remains partially associated, also known as “bearding” (26). The percentages shown are based on analyses of at least 50 cells for each strain with visible signs of sporulation from a single biological replicate.

### SipL’s LysM domain mediates SipL binding to SpoIVA during *C. difficile* sporulation

Since we previously showed that SipL’s C-terminal LysM domain is required for SpoIVA to bind to SipL using recombinant proteins produced in *E. coli* (25, 31), we next wanted to confirm that the LysM domain mediates binding to SpoIVA in *C. difficile*. To this end, we compared the ability of C-terminally FLAG-tagged SipL and SipL_ΔLysM_ to co-immunoprecipitate SpoIVA in sporulating *C. difficile* lysates. In particular, we complemented a Δ*sipL* strain with constructs encoding either wild-type SipL or SipL_ΔLysM_ carrying C-terminal FLAG epitope-tags (3xFLAG). FLAG-tagged wild-type SipL readily pulled-down SpoIVA from sporulating cell lysates, whereas FLAG-tagged SipL_ΔLysM_ failed to pull-down SpoIVA (**Fig. 3**). SpoIVA also did not co-immunoprecipitate with untagged SipL variants. Importantly, FLAG-tagged SipL fully restored functional spore formation to the Δ*sipL* background, whereas the FLAG-tagged SipL_ΔLysM_ failed to complement the Δ*sipL* strain (**Fig. S1**). Taken together, these analyses indicate that SipL binding to SpoIVA depends on SipL’s LysM domain in *C. difficile*, and this interaction appears essential for functional spore formation.

**Figure 3.**
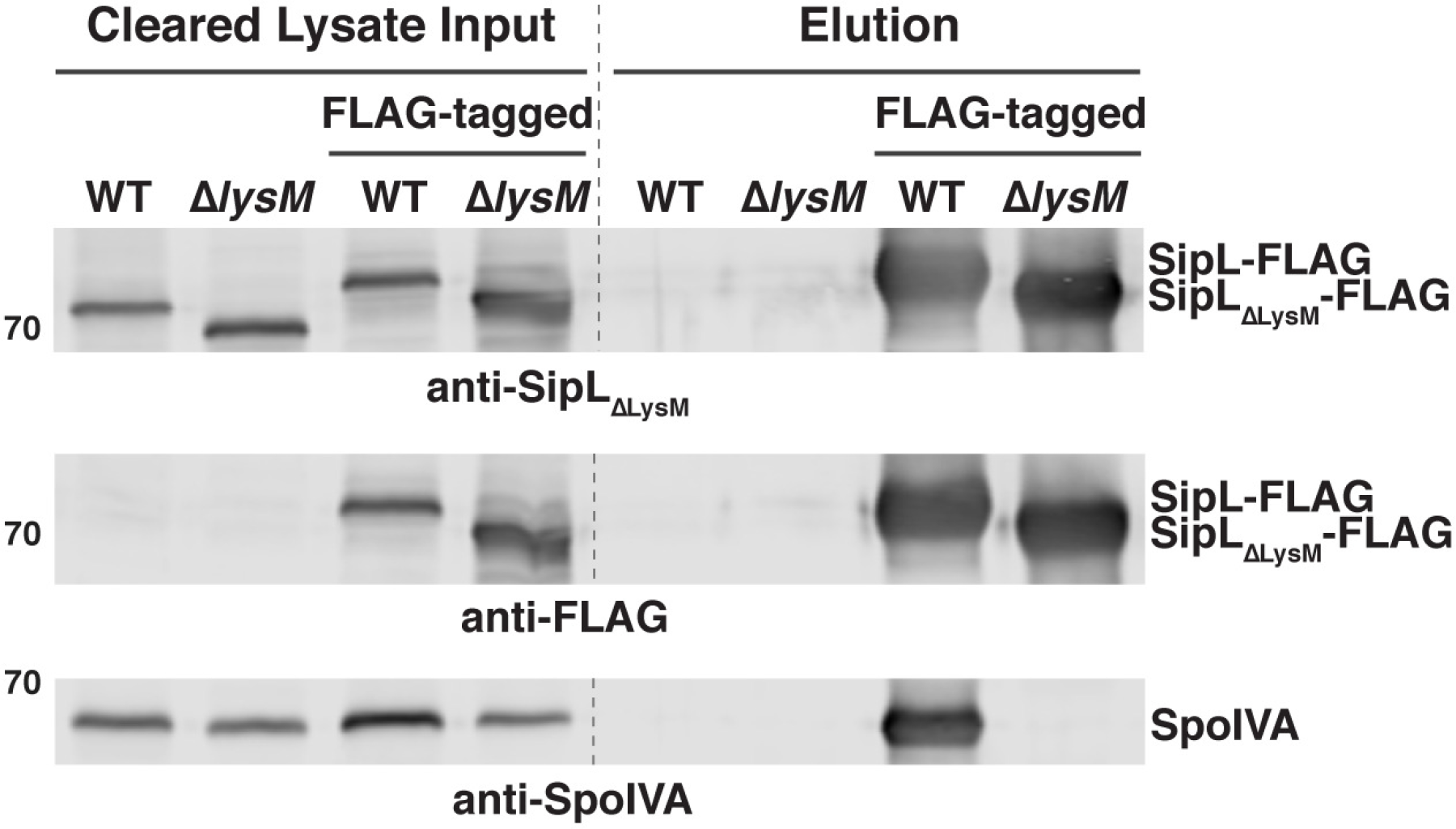
The LysM domain is required for SipL to bind SpoIVA in sporulating *C. difficile* cultures. FLAG-tagged SipL was immunoprecipitated from cleared lysates prepared from the indicated *C. difficile* Δ*sipL* complementation cultures (“Input” fraction) using anti-FLAG magnetic beads. Proteins bound to the beads after several washes were eluted using FLAG peptide (“Elution” fraction). “WT” indicates that Δ*sipL* was complemented with the wild-type *sipL* allele, whereas “Δ*lysM*” indicates that Δ*sipL* was complemented with a *sipL*_Δ*lysM*_ construct. “FLAG-tagged” indicates that the complementation constructs encoded a C-terminal FLAG tag consisting of three successive FLAG tags, which resulted in the SipL-FLAG fusions exhibiting a higher mobility during SDS-PAGE. The untagged *sipL* complementation strains served as negative controls to ensure that untagged SipL and SpoIVA were not non-specifically pulled-down by the anti-FLAG beads. Sporulation was induced for 24 hrs before lysates were prepared. The immunoprecipitations shown are representative of three independent biological replicates.

### Clostridial SipL_LysM_ domains can functionally substitute for the *C. difficile* SipL_LysM_ domain

We next sought to identify key residues within SipL’s LysM domain that are required for the SpoIVA-SipL interaction and thus SipL function. To facilitate the identification of these residues, we tested whether LysM domains from closely related clostridial SipL homologs and SipL’s functional homolog, *B. subtilis* SpoVID (17, 25, 32), could replace the *C. difficile* SipL_LysM_ domain (**Fig. 4A**). Constructs encoding LysM domain chimeras from *Paraclostridium sordellii* and *Paraclostridium bifermentans* SipL homologs (~60% identity and 80% similarity with the *C. difficile* SipL_LysM_ domain), *Clostridium perfringens* SipL (39% identity and 57% similarity), and *B. subtilis* SpoVID (23% identity but 55% similarity) were constructed and used to complement a *C. difficile* Δ*sipL* strain. *P. sordellii* and *P. bifermentans* are members of the same Peptostreptococcaceae family as *C. difficile* (33) but part of a different genus (34), while *C. perfringens* is part of the *Clostridium* genus in the Clostridiaceae family.

**Figure 4.**
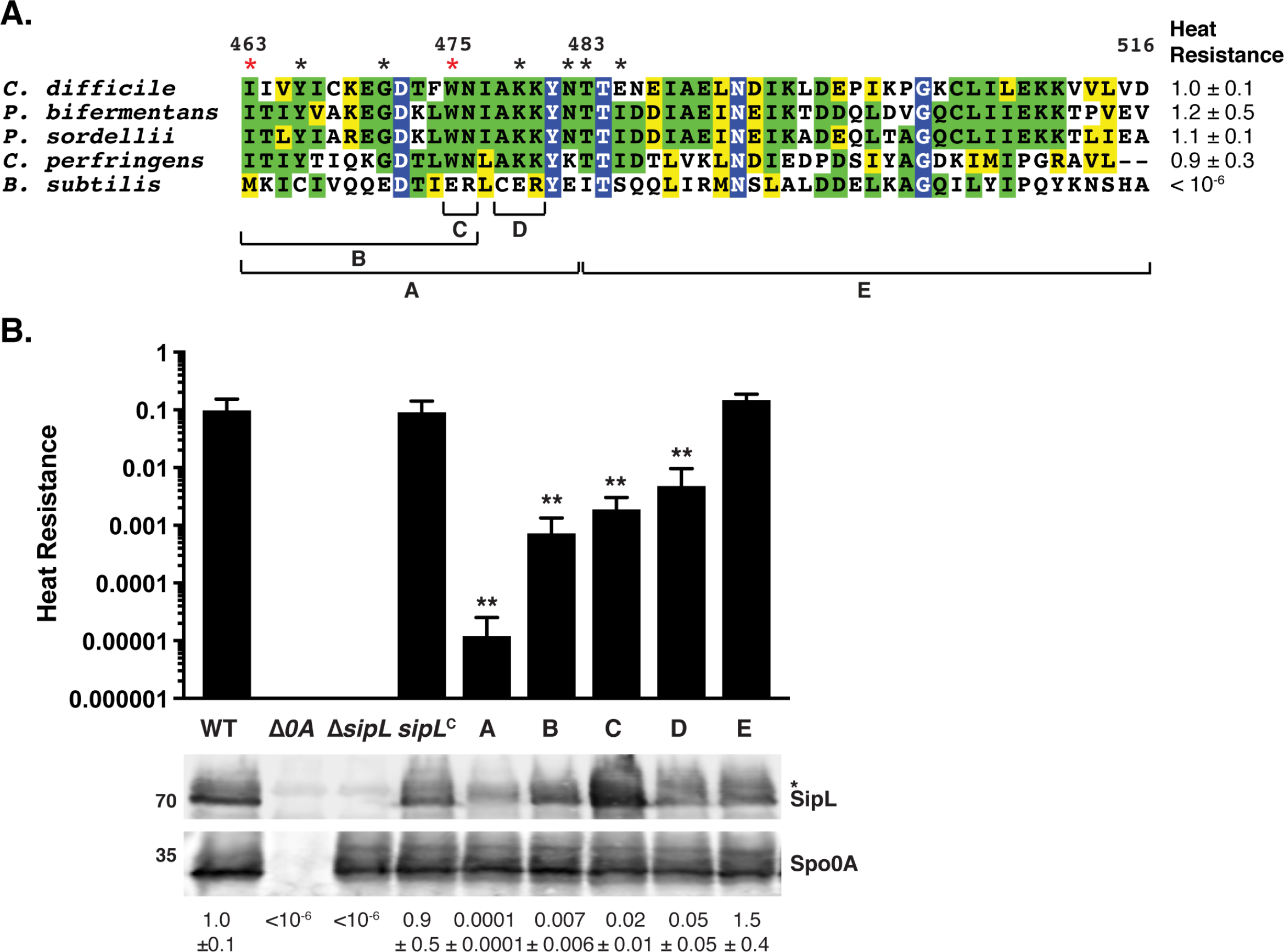
Chimeric analyses identify a small region of the LysM domain important for SipL function. (A) Alignment of LysM domains from SipL homologs from *C. difficile* (CAJ70473), *P. bifermentans* (EQK49575)*, P. sordellii* (EPZ54296), and *C. perfringens* (YP_696893), as well as the LysM domain from *B. subtilis* SpoVID (NP_390689). Blue boxes with white text indicate residues that are completely conserved; green boxes indicate residues that are conserved in some of the homologs; and yellow boxes mark residues that are similar between the homologs. The brackets and letters below the alignment indicate the regions swapped between the *C. difficile* SipL and *B. subtilis* SpoVID LysM domains. The red asterisks highlight residues whose mutations impaired functional spore formation, while black asterisks indicate residues whose mutation to the residue in the *B. subtilis* SpoVID LysM domain did not reduce SipL function, i.e. spore formation (**Fig. S3**). (B) Graphical representation of the heat resistance assay results and western blot analyses of SipL in the chimeric constructs in (A) using an antibody raised against SipL lacking the LysM domain. *sipL*^C^ refers to the Δ*sipL/sipL* wild-type complementation strain. Asterisk marks a non-specific band detected by the anti-SipL_ΔLysM_ antibody. Spo0A was used as a proxy for measuring sporulation induction (4, 25). The western blots shown are representative of the results of three independent biological replicates. The heat resistance efficiencies for all chimeric swaps were determined from 20-24 hr sporulating cultures and represent the mean and standard deviation for a given strain relative to wild type based on a minimum of three independent biological replicates. Statistical significance relative to wild type was determined using a one-way ANOVA and Tukey’s test. ** p < 0.01.

Δ*sipL* strains complemented with the clostridial chimeras (Δ*sipL/sipL-lysM*_*bif*_, Δ*sipL/sipL-lysM*_*sor*_, and Δ*sipL/sipL-lysM*_*per*_) were indistinguishable from wild type and the wild-type *sipL* complementation strain (Δ*sipL/sipL*) by phase-contrast microscopy, since a mixture of phase-bright spores (blue arrows), phase-bright (yellow arrows), and phase-gray (green arrows) forespores were visible in all strains producing clostridial LysM domains (**Fig. S2A**). In contrast, the *B. subtilis* chimera resembled the Δ*sipL/sipL*_Δ*lysM*_ strain, with no phase-bright spores being detected and most of the forespores being phase-dark or phase-gray sporelets (orange arrows). Similar to the parental Δ*sipL* strain, mislocalized coat was visible in the cytosol of the *B. subtilis* LysM chimera strain (pink arrows, **Fig. S2A**).

Consistent with these observations, the clostridial SipL_LysM_ domain swap constructs fully complemented the Δ*sipL* strain in heat resistance assays (**Figs. 4A**and **S2A**), whereas the *B. subtilis* LysM chimeric strain failed to produce heat-resistant spores like the parental Δ*sipL* strain. Importantly, western blotting revealed that the different chimeric SipL variants were produced at relatively similar levels as *C. difficile* SipL using an antibody raised against SipL_ΔLysM_ (**Fig. S2B**), although slightly lower levels of SipL carrying the *B. subtilis* SpoVID LysM domain were observed. Taken together, these results suggest that clostridial SipL_LysM_ domains can still bind *C. difficile* SpoIVA, whereas the *B. subtilis* SpoVID_LysM_ domain cannot.

### Identification of *C. difficile* LysM domain residues important for SipL function

We next used these observations to guide finer-scale chimeric analyses of the *C. difficile* LysM domain by identifying differences between the sequences of the clostridial SipL_LysM_ domains and the *B. subtilis* SpoVID LysM domain. Clostridial LysM domains exhibited the greatest sequence similarity in the N-terminal portion of the LysM domain, so we swapped the residues corresponding to 463-482 of *C. difficile*’s SipL_LysM_ domain with those of the *B. subtilis* SpoIVD LysM domain (Region A, **Fig. 4A**). We also assessed the importance of the C-terminus of the LysM domain by swapping out residues 483-516 (Region E). While the C-terminal chimeric construct, *sipL*_483-516_, fully complemented the Δ*sipL* strain, the *sipL*_463-482_ chimeric construct resulted in an ~3000-fold decrease in heat resistance efficiency relative to wild type and the wild-type *sipL* complementation construct (**Fig. 4B**, p < 0.001). Given the apparent importance of region A, we generated constructs encoding smaller scale swaps, namely residues 463-476 (Region B), 475-476 (Region C), and 478-480 (Region D). Swapping the residues in Regions B and C resulted in an ~100-fold decrease in heat resistance efficiency relative to wild type, while the Region D swap construct resulted in an ~20-fold decrease relative to wild-type (**Fig. 4B**, p < 0.01). Importantly, all the SipL chimeric variants generated produced SipL at wild-type or close to wild-type levels (**Fig. 4B**), consistent with our earlier analyses of the Δ*lysM* strain (**Fig. 1C**).

Based on these findings, we next constructed individual point mutations in Regions C and Region C comprises residues Trp475 and Asn476 in *C. difficile* SipL, which are glutamate and arginine residues, respectively, in the LysM domain of *B. subtilis* SpoVID. We mutated Trp475 to glutamate, since these residues differ more in size and charge than Asn476 relative to arginine. Expression of the *sipL*_W475E_ in Δ*sipL* resulted in a similar ~100-fold defect in heat-resistant spore formation (**Fig. 5A**) relative to wild type as the *sipL*_475-476_ Region B allele, suggesting that the substitution of Trp475 for Glu (W475E) is the major contributor to the sporulation defect of the Region B mutant.

**Figure 5.**
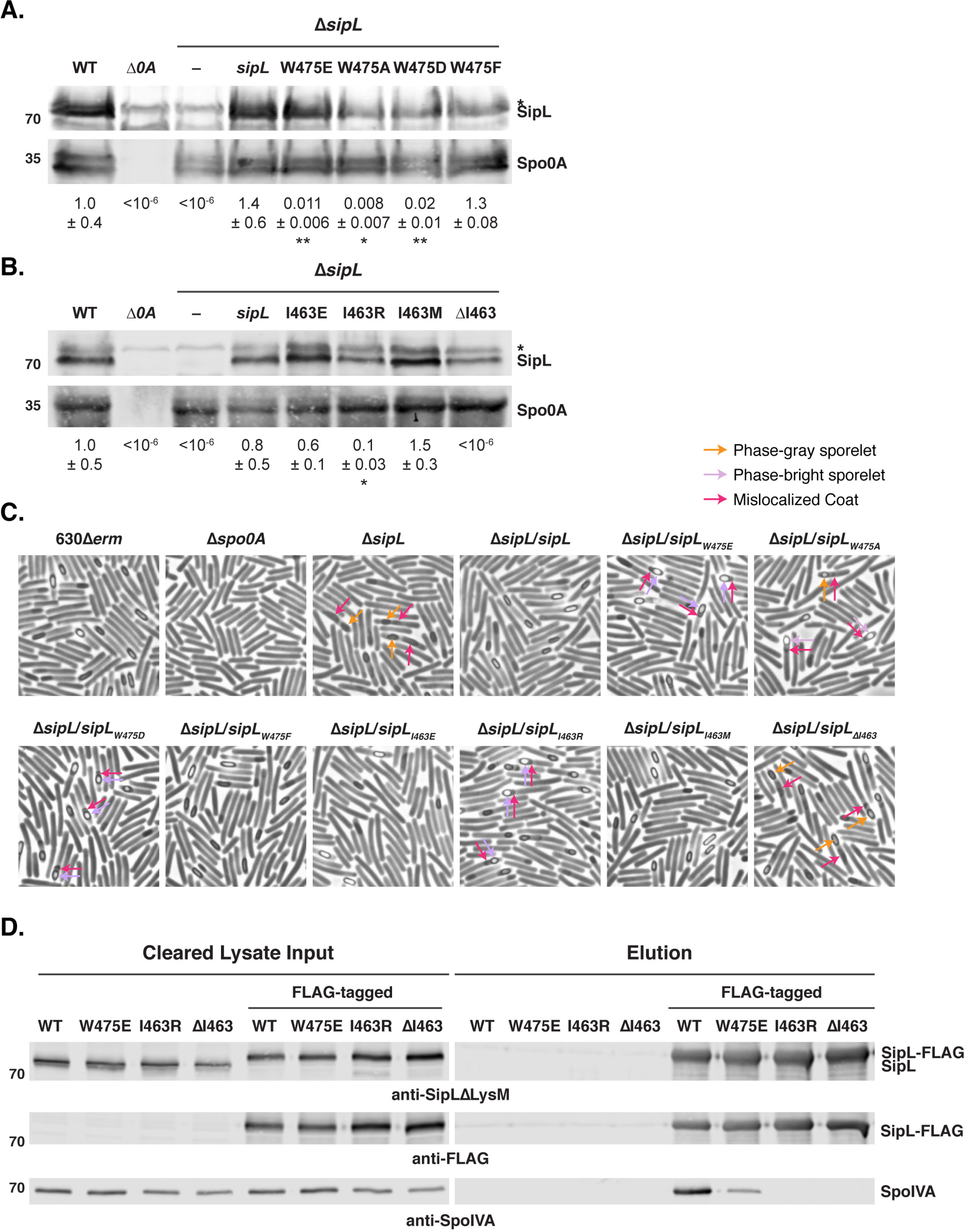
Identification of LysM domain residues required for optimal SipL function. Western blot analyses of strains encoding SipL point mutations in Trp475 (A) and Ile463 (B) using an antibody raised against SipL lacking the LysM domain. Asterisk marks the non-specific band detected by the anti-SipL_ΔLysM_ antibody. Spo0A was used as a proxy for measuring sporulation induction (4, 25). The western blots shown are representative of the results of three independent biological replicates. Heat resistance efficiencies were determined from 20-24 hr sporulating cultures and represent the mean and standard deviation for a given strain relative to wild type based on a minimum of three independent biological replicates. The limit of detection of the assay is 10^-6^. Statistical significance relative to wild type was determined using a one-way ANOVA and Tukey’s test. ** p < 0.01, * p < 0.05. (C) Phase-contrast microscopy analyses of the indicated *C. difficile* strains ~20 hrs after sporulation induction. Δ*sipL* was complemented with either the wild-type allele or the indicated LysM domain mutations. Arrows mark examples of sporulating cells with various defects in maturation: orange arrows highlight phase-gray sporelets (20), which produce a phase-dark ring surrounding the forespore but do not become phase-bright; purple arrows highlight phase-bright sporelets that appear swollen but are outlined by a phase-dark ring; and pink arrows demarcate regions likely to be mislocalized coat (26, 30). The images shown derive from two separate experiments, with the Trp475 and Ile463 variants being performed on different days. (D) Co-immunoprecipitations of FLAG-tagged SipL variants carrying either point mutations in the LysM domain (W475E or I463R) or lacking residue Ile463 (ΔI463). Strains containing untagged SipL were used as negative controls to assess the specificity of the SpoIVA co-immunoprecipitations.

By phase-contrast microscopy, the *sipL*_W475E_ strain resembled the Δ*sipL* strain in producing cells with mislocalized coat (**Fig. 5C**). However, the coat did not appear to be as detached from the forespore of W475E mutant cells relative to the parental Δ*sipL* strain (pink arrows, **Fig. 5C**). Furthermore, W475E produced phase-bright sporelets (purple arrows) as opposed to the phase-gray sporelets (orange arrows) of the parental Δ*sipL* strain.

To further assess the importance of the Trp475 residue for *C. difficile* SipL function, we mutated this residue to the small, uncharged residue, alanine (W475A), and the neutral, aromatic residue, phenylalanine (W475F). We also tested whether reducing the size of the negatively-charged residue by introducing an aspartate at residue 475 instead of a glutamate would affect SipL function. While the conservative change, W475F, did not reduce SipL function, both the W475A and W475D mutations decreased heat-resistant spore formation to levels similar to the W475E mutant (**Fig. 5A**). Since these strains also produced phase-bright sporelets and partially displaced coat (**Fig. 5C∫**like the W475E mutant, these results suggest that the aromatic nature of Trp475 is critical to its function.

We next tested whether a single point mutation in region D, which consists of three residues from 478 to 480, could recapitulate the ~20-fold decrease in heat resistance observed for the region D swap mutant relative to wild type (**Fig. 4**). We focused on lysine 479 in *C. difficile* LysM, since this residue is a lysine in clostridial SipL_LysM_ domains but a negatively-charged glutamate residue in the *B. subtilis* SpoVID LysM domain, and the other two residues in this region carried more conservative changes of alanine 478 to cysteine and lysine 480 to arginine. Substitution of lysine 479 to glutamate did not impair the ability of this allele to complement Δ*sipL* (K479E, **Fig. S3**), suggesting that the other changes in region D could be responsible for the ~20-fold defect observed with this chimeric mutant. However, since the defect was relatively mild relative to region B, we did not further analyze point mutants in this region.

To assess whether additional residues in Region B (463-482) might also contribute to SipL function, we constructed the following point mutant alleles based on differences between the LysM domains of *C. difficile* SipL and *B. subtilis* SpoVID: Y466C, G471E, and N482E. We also tested a few other residues just outside Region B: T483I and E485S. These mutant constructs all fully complemented Δ*sipL* (**Fig. S3**), suggesting that these individual residues do not affect SipL binding to SpoIVA.

We next analyzed the contribution of the first residue of *C. difficile*’s SipL_LysM_ domain because two observations suggested that Ile463 may be important for SipL binding to SpoIVA. Although SipL’s LysM domain is annotated in the NCBI as spanning residues 464-508, we previously observed that a His-tagged construct encoding residues 464 to 516 failed to bind SpoIVA in co-affinity purification analyses in *E. coli*, whereas a construct spanning residues 463-516 resulted in robust pull-down (25). Furthermore, when we tested LysM chimeras using the NCBI-annotated LysM domains of clostridial SipL homologs, all these constructs failed to complement Δ*sipL* (data not shown). However, inclusion of the equivalent Ile463 residue allowed for full complementation (**Fig. 4A**). Taken together, these observations strongly suggest that Ile463 is important for SipL function.

Since Ile463 is a methionine in *B. subtilis* SpoVID, we tested whether the identity and/or precise position of residue 463 is critical for SipL function. Specifically, we generated complementation constructs in which Ile463 was either deleted (ΔI463) or mutated to *B. subtilis* SpoVID’s methionine (I463M), negatively-charged glutamate (I463E), or positively-charged argarginine (I463R). No heat-resistant spores were detected when Ile463 was deleted (**Fig. 5B**), consistent with our observations with the chimeric LysM constructs (data not shown). In contrast, *sipL*_I463M_, the *B. subtilis* SpoVID LysM point mutation, fully complemented the Δ*sipL* mutant (**Fig. 5B**). These results indicate that the sequence difference at residue 463 between the *C. difficile* and *B. subtilis* LysM domains is not responsible for the failure of the SpoVID LysM domain to complement for SipL function. Similarly, only a slight decrease in heat-resistant spore formation was observed with the *sipL*_I463E_ mutation (**Fig. 5B**). In contrast, the *sipL*_I463R_ mutation decreased spore formation by ~9-fold (p < 0.03, **Fig. 5B**), indicating that a positively charged residue at position 463 can impair SipL function.

Phase-contrast microscopy analyses revealed that the I463R mutant produced phase-bright sporelets and partially displaced coat similar to the W475 mutants (**Fig. 5C**), a phenotype that appears to be less severe than loss of SipL or the LysM domain altogether (**Figs. 1B** and **S2**). Indeed, the sipL_ΔI463_ strain resembled the parental Δ*sipL* strain in that only phase-gray sporelets (purple arrows, **Fig. 5C**) were observed, and the coat displacement appeared more severe than the W475 and I463R point mutant strains (pink arrows). Taken together, our results suggest that the spacing of the LysM domain, i.e. starting at position 463, is critical for SipL function, while the chemical identity of the residue at this position can also impact SipL function.

### Isoleucine 463 and Tryptophan 475 enhance binding of SipL to SpoIVA

The decreased ability of *sipL* constructs encoding mutations in either Ile463 or Trp475 to complement Δ*sipL* implied that point mutations in these residues decrease SipL binding to SpoIVA. To test this hypothesis, we generated constructs encoding FLAG-tagged SipL that carry mutations in these residues and measured their ability to pull-down SpoIVA in co-immunoprecipitation analyses. Whereas SpoIVA was readily pulled-down by FLAG-tagged wild-type SipL, no SpoIVA was detected in immunoprecipitations with SipL variants where Ile463 was either deleted (ΔI463) or mutated to arginine (I463R, **Fig. 5D**). Reduced amounts of SpoIVA were pulled down with the FLAG-tagged SipL_W475E_ variant relative to wild type. Interestingly, the pull-down results did not entirely match the severity of the mutations, since the *sipL*_W475E_ allele resulted in an ~100-fold heat resistance defect, but the *sipL*_I463R_ allele caused only a ~9-fold heat resistance defect. This result suggests that even though SpoIVA can bind SipL_W475E_, the interaction does not result in optimal SpoIVA and/or SipL function.

### SipL localization to the forespore depends on the LysM domain

Since SpoIVA binding to SipL’s LysM domain appears critical for these proteins to mediate spore formation, we hypothesized that SipL’s LysM domain would be important for recruiting SipL to the forespore and allowing it to encase the forespore. To directly test this hypothesis, we complemented Δ*sipL* with a construct encoding an mCherry fusion to SipL_ΔLysM_. Whereas wild-type SipL fused to mCherry encases the forespore (**Fig. 6A**) as previously reported (27), deletion of the LysM domain prevented SipL_ΔLysM_-mCherry localization to the forespore, since the SipL_ΔLysM_-mCherry signal was largely distributed in the mother cell cytosol (**Fig. 6A**). Western blot analyses revealed that mCherry was not liberated from SipL_ΔLysM_-mCherry (**Fig S4A**), indicating that SipL’s LysM domain directs SipL to the forespore and allows it to encase the forespore, presumably through its interaction with SpoIVA.

**Figure 6.**
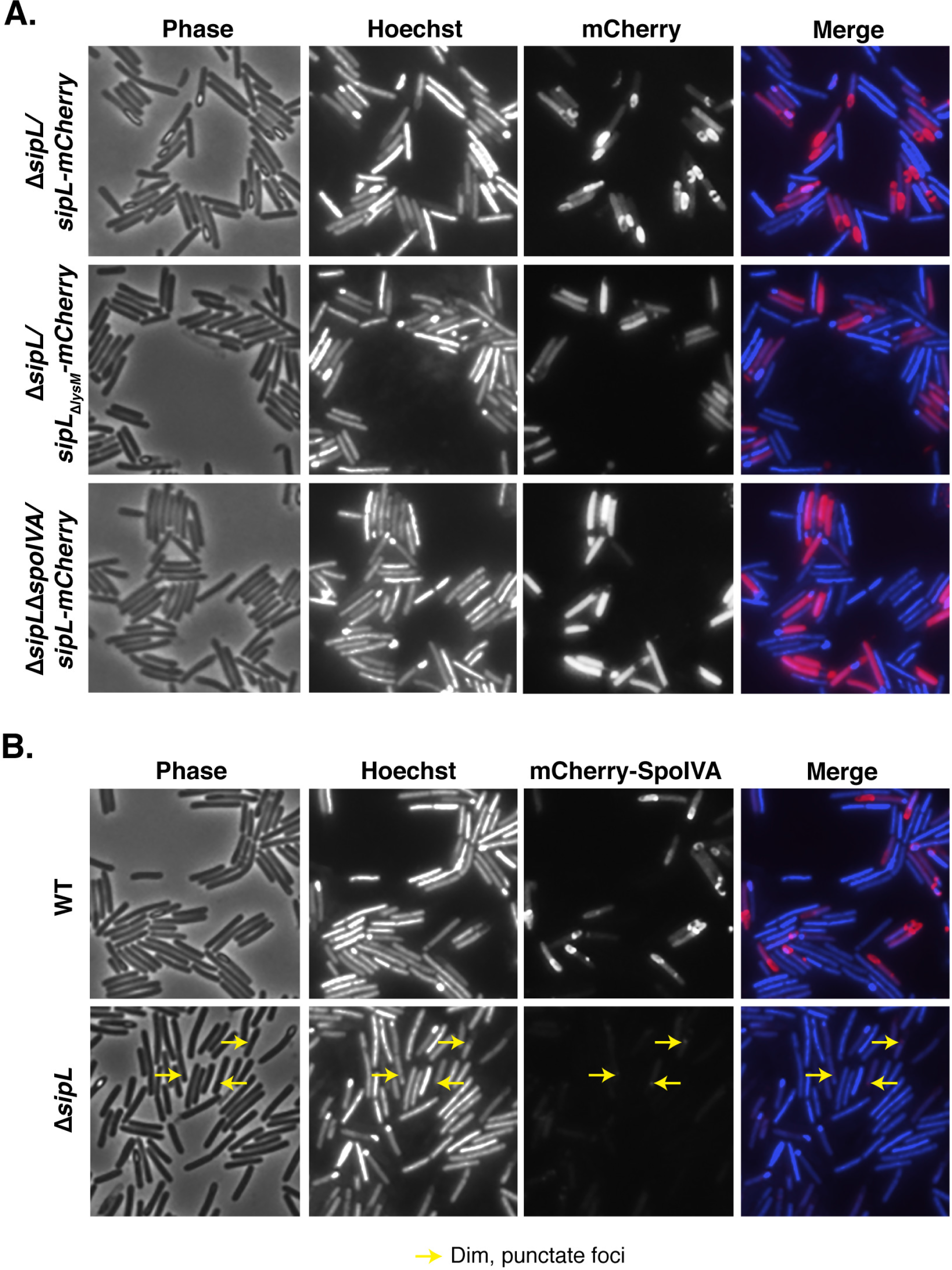
SipL binding to SpoIVA is required to localize SipL to the forespore, while SpoIVA encasement of the forespore depends on SipL. (A) Fluorescence microscopy analyses of either Δ*sipL* or Δ*sipL*Δ*spoIVA* complemented with constructs encoding either SipL-mCherry or SipL_ΔLysM_-mCherry after 20-23 hrs post-sporulation induction. (B) mCherry-SpoIVA localization in either the wild-type or Δ*sipL* strain backgrounds. A wild-type copy of *spoIVA* was present in both strains because the fusion protein does not efficiently encase the forespore unless a wildtype copy of *spoIVA* is present (26). Cells were visualized using phase-contrast (phase) microscopy, and the nucleoid was visualized using Hoechst staining (41). The Hoechst-stained nucleoid is shown in blue, and mCherry fluorescence is shown in red. Engulfment completion excludes Hoechst from staining the forespore (41). The merge of Hoechst and mCherry is also shown. Yellow arrows mark single, dim foci of mCherry-SpoIVA observed in the Δ*sipL* strain background. The images shown are representative of three independent biological replicates.

### SipL requires SpoIVA to localize to and encase the forespore and encase it, while SpoIVA requires SipL to encase the forespore

To directly assess whether SpoIVA was required for SipL to localize to the forespore, we analyzed the localization of SipL-mCherry in a Δ*spoIVA*Δ*sipL* mutant. It was necessary to use the double mutant because the presence of wild-type (untagged) SipL results in some SipL-mCherry being re-distributed to the cytosol (data not shown). In the absence of SpoIVA, the SipL-mCherry signal was entirely cytosolic similar to the localization pattern of SipL_ΔLysM_-mCherry (**Fig. 6A**), although the cytosolic SipL-mCherry signal appeared more intense in the absence of SpoIVA.

Since these results indicated that SipL localization to and around the forespore depends on SpoIVA through its interaction with SipL’s LysM domain, we next assessed whether SpoIVA’s localization around the forespore (26) depends on SipL. To test this question, we analyzed the localization of mCherry-SpoIVA, which has almost wild-type function (26), in the absence of SipL. For these experiments, mCherry-SpoIVA localization was analyzed in the presence of wild-type SpoIVA because mCherry-SpoIVA exhibits reduced encasement of the forespore if it is the only variant of SpoIVA present (26). While mCherry-SpoIVA encased the forespore (with some cytosolic localization) when produced in a wild-type strain background, mCherry-SpoIVA produced in the absence of SipL localized to a single, albeit dim, focus on the forespore at the mother cell proximal side (**Fig. 6B**). Notably, the amount of mCherry-SpoIVA produced in the Δ*sipL* background was reduced relative to the wild-type background (**Fig. S4**), consistent with our original finding that SpoIVA levels are reduced in the absence of SipL (25). Taken together, our results indicate that SpoIVA-SipL binding is needed to not only bring SipL to the forespore but also for SipL to encase the forespore; in contrast, SpoIVA can find the forespore in the absence of SipL, but SpoIVA requires SipL to surround the forespore.

## Discussion

In this study, we show that the LysM domain of *C. difficile* SipL is critical for functional spore formation because it binds to SpoIVA and allows both these proteins to encase the developing forespore. Our conclusions are based on the following observations: (i) SipL-mCherry mislocalizes to the cytosol if either SpoIVA or SipL’s LysM domain is absent (**Fig. 6**); (ii) deletion of the LysM domain results in coat mislocalization to the mother cell cytosol (**Fig. 2**) and prevents heat-resistant spore formation (**Fig. 1**) analogous to the *sipL* deletion mutant, and SpoIVA cannot bind to SipLΔ_LysM_ in immunoprecipitation analyses (**Fig. 3**); and (iii) specific point mutations in SipL’s LysM domain impair binding to SpoIVA and reduce functional spore-formation (**Fig. 5**).

Through chimeric LysM analyses that exploited the failure of the *B. subtilis* SpoVID LysM domain to complement *C. difficile* LysM domain function (**Fig. 4**), we identified two point mutations in the LysM domain, W457E and I463R, that decrease heat-resistant spore formation by decreasing or even preventing binding to SpoIVA (**Fig. 5**). While these mutations reduced spore formation by ~100-and 10-fold, respectively, their effects on coat mislocalization were qualitatively less severe. The coat in *sipL*_W475E/D/A_ and *sipL*_I463R_ mutants appeared more closely associated with the forespore in these strains, particularly if the forespores/sporelets appeared phase-bright (**Fig. 5C**). In contrast, the coat appeared to be more frequently displaced to the mother cell cytosol in *sipL* mutants carrying non-functional *sipL* alleles, like Δ*lysM*, *sipL*_ΔI463_, and *sipL*-*lysM*_*Bsub*_ (**Figs. 1, 5**, and **S2**). Indeed, the *sipL* strains that failed to produce heat-resistant spores did not make the phase-bright, swollen sporelets observed in *sipL*_W475E/D/A_ and *sipL*_I463R_ mutants (**Fig. 5C**, purple arrows). It would be interesting to test whether the *sipL*_W475E/D/A_ and *sipL*_I463R_ mutants localize coat close to the forespore because SipL_W457E/D/A_ and SipL_I463R_ variant can still localize to the forespore and/or partially encase the forespore. This scenario seems possible for the *sipL*_W457E_ allele, since SipL_W475E_ partially binds SpoIVA (**Fig. 5D**).

Our analyses also implicated the first residue of the LysM domain as being important for SipL function. The I463R mutation, but not the I463M or I463E mutations, significantly impaired SipL function, while deletion of the Ile463 residue altogether completely abrogated SipL function (**Fig. 5**). These observations suggest that Ile463 is needed as a linker between the LysM domain and the rest of the SipL protein, or it could be needed for proper folding of the LysM domain. This latter possibility seems less likely given that Ile463 is not strongly conserved at this position across LysM domains (**Fig. S5**) and does not appear to play an important structural role in the LysM domains whose structures have been solved (35).

LysM domains frequently bind to N-acetylglucosamine (NAG) residues in chitin (e.g. in eukaryotic LysM domains) and peptidoglycan in prokaryotic LysM-containing proteins (35, 36). Recent structural analyses have revealed the molecular basis for LysM binding to peptidoglycan (36). Interestingly, some of the residues identified as being critical for recognizing NAG in an *Enterococcus faecalis* LysM domain are not conserved in the four clostridial SipL_LysM_ domains analyzed in this study (**Fig. S5A**, red asterisks), although they typically have similar properties in clostridial LysM domains. Nevertheless, when the structure of *C. difficile* SipL_LysM_ domain is predicted using the iTasser algorithm (37), it aligns closely with the structures of several LysM domains (**Fig. S5B**), raising the possibility that SipL_LysM_ domains can bind peptidoglycan.

While this possibility remains to be tested, we note that the *B. subtilis* SpoVID_LysM_ domain does not bind peptidoglycan even though putative peptidoglycan-binding residues are conserved (32). In contrast, the LysM domain of SafA, a coat morphogenetic protein that modulates inner coat assembly downstream of SpoVID (38, 39), functions both as a protein-protein interaction module and a peptidoglycan binding domain (32). Specifically, SafA’s LysM domain binds to SpoVID early during coat morphogenesis, while later during morphogenesis, SafA’s LysM domain binds to the spore cortex, the modified peptidoglycan layer that confers heat resistance to spores (32). Thus, even though SafA lacks a transmembrane domain to span the outer forespore membrane, its N-terminal LysM domain apparently binds the spore cortex, since SafA exhibits aberrant localization in cortex biogenesis mutants (32). It remains unclear how SafA binding switches from SpoVID to the cortex during spore formation, but a similar event would need to occur if *C. difficile* SipL’s LysM domain binds SpoIVA and then to the cortex region. Directly testing whether clostridial SipL_LysM_ domains can bind peptidoglycan would provide important insight to these questions.

Interestingly, in the structure model generated by iTasser (**Fig. S5B**), the Trp457 residue we identified as being important for SipL to bind SpoIVA (**Fig. S5B**) is predicted to be surface-exposed. Thus, Trp457 would appear to be available to directly interact with as-yet-undefined regions of SpoIVA and possibly also to bind cortex peptidoglycan. Future studies directed at identifying regions with SpoIVA that mediate binding to SipL will provide important insight into how the interaction between these two proteins allows for functional spore formation. Given that our results indicate that SpoIVA encasement of the forespore (**Fig. 6**) depends on SipL, it is possible that SipL binding to SpoIVA promotes SpoIVA polymerization. Alternatively, reduced levels of mCherry-SpoIVA in the Δ*sipL* strain may prevent self-polymerization of SpoIVA (40) and thus prevent encasement. Determining the precise effects of SipL binding to SpoIVA on SpoIVA’s presumed ATPase and polymerization activities will provide critical insight into how infectious *C. difficile* spores are built and could guide efforts to prevent spore formation by clostridial pathogens.

## Materials and Methods

### Bacterial strains and growth conditions

630Δ*erm*Δ*pyrE* (28) was used as the parental strain for *pyrE-*based allele-coupled exchange (ACE, (28)). *C. difficile* strains are listed in **Table S1** and were grown on BHIS agar (42) supplemented with taurocholate (TA, 0.1% w/v; 1.9 mM), kanamycin (50 μg/mL), and cefoxitin (8 μg/mL) as needed for conjugations. *C. difficile* defined media (CDDM, (43)) was used for isolating complementation strains (28). 5-fluoroorotic acid (5-FOA) at 2 mg/mL and uracil at 5 μg/mL as needed for ACE. Cultures were grown under anaerobic conditions using a gas mixture containing 85% N_2_, 5% CO_2_, and 10% H_2_.

*Escherichia coli* strains for HB101/pRK24-based conjugations and BL21(DE3)-based protein production are listed in **Table S1**. *E. coli* strains were grown at 37°C, shaking at 225 rpm in Luria-Bertani broth (LB). The media was supplemented with chloramphenicol (20 μg/mL) and ampicillin (50 μg/mL) as needed.

### *E. coli* strain construction

All primers are listed in **Table S2**, and all g-blocks used for cloning are listed in **Table S3**. Details of *E. coli* strain construction are provided in the **Supplementary Text S1**. All plasmid constructs were cloned into DH5α and sequenced confirmed using Genewiz. Plasmids to be conjugated into *C. difficile* were transformed into HB101/pRK24, while the plasmid used for antibody production was transformed into BL21(DE3).

### *C. difficile* strain complementation

Allele-coupled exchange (ACE, (28)) was used as previously described (44) to construct the Δ*spoIVA*Δ*sipL*Δ*pyrE* double mutant. Δ*spoIVA*Δ*pyrE* was used as the parental strain, with strain #1704 pMTL-YN3 Δ*sipL* being used to introduce the *sipL* mutation. pMTL-YN1C-based complementation constructs were conjugated into Δ*sipL*Δ*pyrE* as previously described (44). Two independent clones of each complementation strain were phenotypically characterized.

### Plate-based Sporulation

*C. difficile* strains were grown from glycerol stocks overnight on BHIS plates containing TA (0.1% w/v). Colonies from these cultures were then used to inoculate liquid BHIS cultures, which were grown to stationary phase and then back-diluted 1:50 into BHIS. When the cultures reached an OD_600_ between 0.35 and 0.7, 120 μL was removed to inoculate 70:30 agar plates ((25)). Sporulation was induced on this media for 20-24 hrs. The ~20 hr timepoint was used to analyze cultures by phase-contrast microscopy and harvest samples for Western blot analyses.

### Heat resistance assay on sporulating cells

Heat-resistant spore formation was measured in sporulating *C. difficile* cultures after 20-24 hrs as previously described (45). Briefly, sporulating cultures were divided into two, with one culture being heat-treated at 60°C for 30 min, while the second half was left untreated. The samples were serially diluted and plated on BHIS(TA), and the heat resistance (H.R.) efficiency calculated from the colonies that arose. Specifically, the H.R. efficiency represents the average ratio of heat-resistant cells for a given strain relative to the average ratio determined for wild type based on a minimum of three biological replicates. Statistical significance was determined using a one-way ANOVA and Tukey’s test.

### Antibody production

The anti-SipL_ΔLysM_ antibody used in this study was raised against SipL_ΔLysM_-His_6_ in a rabbit by Cocalico Biologicals (Reamstown, PA). The recombinant protein was purified from *E. coli* strain #764 (25) (**Table S1**) using Ni^2+^-affinity resin as previously described (46).

### TEM analyses

Sporulating cultures (23 hrs) were fixed and processed for electron microscopy by the University of Vermont Microscopy Center as previously described (25). A minimum of 50 full-length sporulating cells were used for phenotype counting.

### Immunoprecipitation analyses

Sporulation was induced on 70:30 plates for 24 hrs as described above. Cultures from three 70:30 plates per strain were scraped into 3 x 1 mL of FLAG IP buffer (FIB: 50 mM Tris pH 7.5, 150 mM NaCl, 0.02% sodium azide, 1X Halt protease inhibitor (ThermoScientific)). The cultures were transferred into tubes containing a ~300 μL of 0.1 mm zirconia/silica beads (BioSpec). The cultures were lysed using a FastPrep-24 (MP Biomedicals) for 4 x 60 s at 5.5 M/s, with 5 min cooling on ice between lysis intervals. The tubes were pelleted at 14,500 g for 10 min at 4°C to pellet beads, unbroken cells/spores, and insoluble material. After washing the tubes with additional FIB, the lysates were pooled and FLAG-conjugated magnetic resin was added. This magnetic resin was generated by incubating Dynabead Protein G (ThermoScientific) with anti-FLAG antibodies (Sigma Aldrich) at room temperature followed by washing. The lysates were incubated with the anti-FLAG resin for 2 hrs at room temperature with rotation. After washing the beads with FIB, FLAG-tagged proteins were eluted using FIB containing 0.1 mg/mL FLAG peptide (Sigma Aldrich). All immunoprecipitations were performed on three independent biological replicates.

### mCherry fluorescence microscopy

Live cell fluorescence microscopy was performed using Hoechst 33342 (Molecular Probes, 15 μg/mL) and mCherry protein fusions. Samples were prepared on agarose pads as previously described (30), and samples were imaged 30 min after harvesting to allow for mCherry fluorescence signal reconstitution in the anaerobically grown bacteria as previously described (27). Briefly, phase-contrast and fluorescence microscopy were performed using a Nikon 60x oil immersion objective (1.4 NA) on a Nikon 90i epifluorescence microscope. A CoolSnap HQ camera (Photometrics) was used to acquire multiple fields for each sample in 12 bit format using NIS-Elements software (Nikon). The Texas Red channel was used to acquire images after a 3-90 ms exposure (90 ms for SipL-mCherry and mCherry-IVA), 50 ms for Hoechst staining, and ~3 ms exposures for phase-contrast microscopy) with 2 x 2 binning.

Ten Mhz images were subsequently imported into Adobe Photoshop CC 2015 for minimal adjustments in brightness/contrast levels and pseudocoloring. Localization analyses were performed on three independent biological replicates.

### Western blot analyses

Samples for western blotting were prepared as previously described (25). Briefly, sporulating cell pellets were resuspended in 100 μL of PBS, and 50 μL samples were freeze-thawed for three cycles and then resuspended in 100 μL EBB buffer (8 M urea, 2 M thiourea, 4% (w/v) SDS, 2% (v/v) β-mercaptoethanol). The samples were boiled for 20 min, pelleted, re-suspended in the same buffer to maximize protein solubilization, boiled for another 5 min and then pelleted. Samples were resolved on 12% SDS-PAGE gels, transferred to Immobilon-FL PVDF membrane, blocked in Odyssey^®^ Blocking Buffer with 0.1% (v/v) Tween 20. Rabbit anti-SipL_ΔLysM_ and mouse anti-SpoIVA (47) were used at 1:2,500 dilutions; rabbit anti-mCherry (Abcam) was used at a 1:2,000 dilution; and rabbit or mouse anti-Spo0A (25, 31) was used at a 1:1,000 dilution. IRDye 680CW and 800CW infrared dye-conjugated secondary antibodies were used at a minimum of 1:25,000 dilution, and blots were imaged on an Odyssey LiCor CLx. Western blots were performed on sporulating samples derived from three independent biological replicates.

## Acknowledgments

We would like to thank N. Bishop and J. Schwarz for excellent assistance in preparing samples for transmission electron microscopy throughout this study; D. Weiss and C. Ellermeier for providing the codon-optimized mCherry construct (48); A. Camilli for access to the Nikon microscope; N. Minton (U. Nottingham) for generously providing us with access to the 630Δ*erm*Δ*pyrE* strain and pMTL-YN1C and pMTL-YN3 plasmids for allele-coupled exchange (ACE); M. Dembek for directly providing these materials to us and sharing his specific protocols on ACE.

Research in this manuscript was funded by Award Number T32AI007329 to M.H.T. and R01AI22232 from the National Institutes of Allergy and Infectious Disease (NIAID) to A.S. A.S. is a Pew Scholar in the Biomedical Sciences supported by The Pew Charitable Trusts and a Burroughs Wellcome Investigator in the Pathogenesis of Infectious Disease supported by the Burroughs Wellcome Fund. The content is solely the responsibility of the author(s) and does not necessarily reflect the views of the Pew Charitable Trusts, Burroughs Wellcome, NIAID, or the National Institutes of Health. The funders had no role in study design, data collection and interpretation, or the decision to submit the work for publication.

A.S. has a paid consultancy for BioVector, Inc., a diagnostic start-up.

